# Effect of Nitrogen Fertilizer Rate on Yield and Quality Parameters of Malt Barley Varieties for Beer Production in South Gonder, Ethiopia

**DOI:** 10.1101/2022.09.12.507533

**Authors:** Admas Alemu Abebe, Fikre Hagos

## Abstract

The objective of this study was to examine the effect of different nitrogen rates on the yield and malt barley quality parameters of two malting barley varieties (Holker and miskal-21). The experiment was carried out during 2013 cropping season at three locations, namely Ebnat, Estie and Farta. The following nitrogen rates were applied: 0, 60, 120and 180 kg ha-1 and a factorial experiment arranged in RCBD was applied to conduct the research. The result of this study indicated that nitrogen significantly increased the grain yield (648.2 to 1657.4and 666.7 to 1851.8 kg ha^-1^for Holker and Miskal-21 respectively). Malt quality was decreased by increasing nitrogen rates (1.28 to 2.34%). Apart from nitrogen rates, the varieties also affected the yield and nature of the protein content levels. As indicated in this result Holker variety gave optimum grain yield and lowest protein content, therefore, this variety is advisable to be used by malt brewers to satisfy their requirements. The optimal malt quality and yield was achieved by applying between 60-120 kg N ha-1 beyond this rate, grain yield might be significantly increased but malt qualities at the same time significantly decreased therefore it is recommended to obtain optimum yield as well as high malt qualities

## 1. INTRODUCTION

Barley (*Hordeum vulgare* L.) is an important crop in the world mainly used for animal feed and malt. New winter barley genotypes have a high yield potential, nearly as the winter wheat. Agro-ecological conditions and production technology, primarily the natural soil fertility, the amount of applied nitrogen (Abeledo *et al*., 2003), and primarily the carbon to nitrogen (C/N) ratio (Acuna *et al*., 2005), significantly affected the grain yield of malting barley. Leistrumaite (2005) emphasized that the grain yield and quality traits of spring barley varieties varies greatly due to growing conditions. Regarding the barley quality, it was observed that high protein content decreases the extract yield, turbids the beer and slows down the start of germination, while too low protein content results in a lower enzymatic activity and slow growth of yeast in brewery (Pettersson, 2007).

Nitrogen (N) is often the most limiting factor in crop production. Hence, application of fertilizer nitrogen results in higher biomass yields and protein yield and concentration in plant tissue is commonly increased. Nitrogen often affects amino acid composition of protein and in turn its nutritional quality. In cereals, abundant supply of nitrogen decreases the relative proportion of lysine and threonine, thus, reducing the biological value of the protein. Increasing nitrogen supply generally improves kernel integrity and strength, resulting in better milling properties of the grain (Jürg *et al*., 2008).

Cereal breeders select barley for large grain, thin husk and low protein content to improve malt quality, and select barley for high protein content on account of animal feed (Pržulj *et al*., 2010). For production of the European type of beer, the optimal crude protein content (CP) is 10.07%, and the optimal dry matter (DM) content ranges from 9.5 to 11.5% (Palmer, 2000). Malting barley breeders all over the world are challenged by the difficulty in producing barley with appropriate CP level (Wang *et al*., 2001; McKenzie *et al*., 2005; Mengel *et al*., 2006). One reason for difficulties in establishing acceptable CP content of grain is that it depends on the nitrogen (N) supply from fertilizer and soil, as well on the carbon (C) supply from the atmosphere. This renders N and C dynamics in the soil/plant system sensitive to environmental conditions (Pettersson, 2007).

Ethiopia has a shortage of malt barley to meet the demand of the local breweries (Mohammed and Getachew, 2003). Limited tests on grain qualities of malt barley have been carried out since 1982 at Holetta barley quality laboratory (HBQL) (Fekadu et al., 1996). Yield and quality of malt barley are influenced by many factors (Teshome *et al*., 2011). Malting barley is grown as a cash crop. The most popular use of malt is the production of alcoholic beverages, but malt and malt extracts are increasingly becoming very important in the bakery and baby food industry. Globally, the demand for beer is rapidly growing, particularly in many developing countries. This makes malting barley an important cash crop for resource-poor farmers in areas where options are very limited, and barley is often the only possible crop (Feiten et al., 2010).

### Objectives

#### General objective

✓ To evaluate the effect of nitrogen fertilizer on Malt barley yield and quality parameters in relation to Factory quality requirement

#### Specific objectives

✓ To identify the best nitrogen rate for best factory quality for each malt barley variety
✓ To identify the best nitrogen rate for best factory quality for each location of the selected weredas
✓ To identify barley varieties for different end uses

## 2. MATERIALS AND METHODS

### 2.1. Description of the Study Area

The field study was conducted at three Weredas found in South Gondar zone Viz., Ebnat, Farta and East Estie during 2014/15 main rain season. The altitude range of the study areas was from 1900 - 4135, 1500 - 4000 and 1500 - 2150 meters above sea level for Farta, East Estie and Ebinat woredas respectively while the range of annual temperature of Farta, East Estie and Ebinat was 4.9-18.4, 8.3-25 and 23-30 °C respectively. The mean annual rainfall for Farta, East Estie and Ebinat ranged from 900-1400, 1307-1500 and 772-1160 mm (source: SGZARDO).

### 2.2. Material Description

Two malt barley varieties (i.e. Miskal-21 and Holker) that are nationally released so far was used for this study to determine the effect of nitrogen fertilizer on malt barley in relation to factory quality parameters. The seed materials were collected from Holeta agricultural research center.

### 2.3. Experimental Design

Two malt barley varieties (Miskal-21and Holker) and four nitrogen rates (0, 60, 120 and 180 kgN/ha) (Donovan *et al*., 2013) was arranged in a factorial RCBD in 3 replications to determine the effect of nitrogen fertilizer on malt barley quality parameters at three weredas of South Gonder zone. The varieties were planted at the rate of 75 kg/ha in the experimental field with a plot size of 12 m^2^ (4m x 3m), spacing of 0.2 m between rows, 1 m between plots and in each weredas, three farmer’s field was used as replications. Planting was carried out on the beginning of June of 2006 E.C. Nitrogen fertilizer was applied two times; one at planting (0, 20, 40 and 60 kg N/ha respectively for the four nitrogen rates) and the other one month after planting (0, 40, 80 and 120 kg N/ha respectively). Weeds were removed 40 days after planting (DAP) and the second weeding was applied 35 days after the first weeding. Furthermore, standard agronomy practices for malting barley were applied. Sowing was carried out in the beginning of June during 2006 E.C. Necessary crop protection measures was performed. The harvested seeds were taken to laboratory for quality parameter analysis at Bahirdar Universities.

### 2.4. Methods of Data Collection

Data was collected as yield/plot for yield then converted to yield/ha and for the quality parameters (Teshome *et al*., 2011) and grain samples from the harvested malt barley varieties was used to determine the characteristics. Protein content, free Amino Nitrogen (FAN) and moisture contents were analyzed for malt grain quality parameters. Malt barley grain was cleaned through screeners to remove stones, foreign bodies, dust, and straw.

**Free Amino Nitrogen (FAN):** This was analyzed by micro-*Kjeldahl* method using 25 mL wort as described in the AOAC (1990) method 950.10.

**Moisture content of barley:** Moisture content of barley was determined using NIR equipment that has been calibrated by the standard ASBC method (ASBC Barley 5C).

**Protein content (N x 6.25):** Protein content was determined on dockage-free barley using NIR equipment that has been calibrated by Combustion Nitrogen Analysis (CNA). CNA is determined on a LECO Model FP-428 CNA analyser calibrated by EDTA. Samples are ground on a UDY Cyclone Sample Mill fitted with a 1.0-mm screen. A 200-mg sample was analysed as received (it is not dried prior to analysis). A moisture analysis is also performed and results are reported on a dry matter basis (ASBC Barley 7C).

### 2.5. Statistical Analysis

Analysis of variance (ANOVA) for malt quality parameters of the varieties subjected to different nitrogen rates was analyzed using SAS (statistical analysis system) for two factors arranged in a factorial RCBD version 9.2, 1999 to 2000, SAS Institute inc., Cary, NC, USA. Means of each character for each variety was compared by LSD at a probability level of 5%.

## 3. RESULT AND DISCUSSION

### 3.1. Grain Yield

There were significant(P*<*0.001) differences among N fertilizer rates in grain yield (Table 2).N fertilizer had shown significant influence on grain yield of malt barley for the tested three locations as indicated in (Table 2). Our results are in line with AlirezaAlazmani, (2014) who reported that nitrogen application had high effects on grain yield of barley cultivar

**Table 2.** Description of malt barley varieties.

| Variety | Altitude (m) | Rainfall (mm) | Maturation period (day) | Grain yield, q/ha |  |
| --- | --- | --- | --- | --- | --- |
|  |  |  |  | On station | On farm |
| Holker | 2300-3500 | 500-700 | 135-141 | 24-38 | 20-25 |
| Miskal-21 | 1550-2800 | 500-700 | 136 | 19-52 | 20-46 |

Source: SGZARDO, 2004 E.C

**Table 2.**
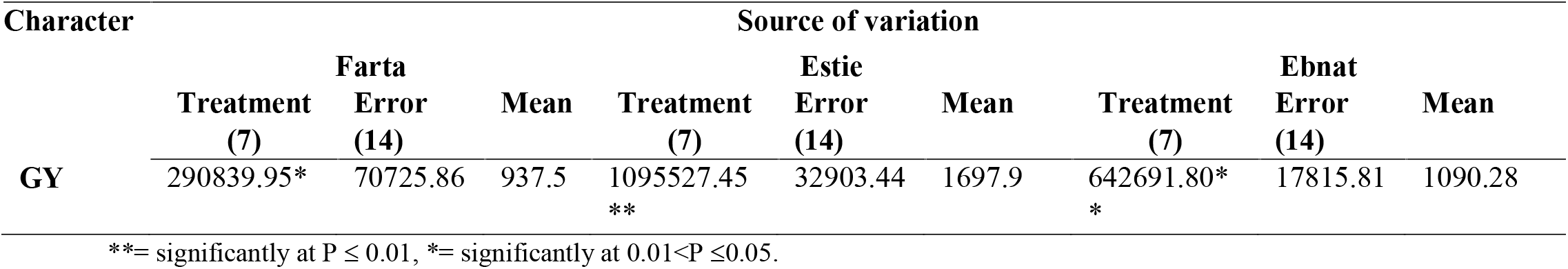
Mean square of characters from separate analysis for grain yield.

The highest grain yield mean recorded from Farta, Estie and Ebnat was 937.50, 1697.92 and 1090.28 kg ha^-1^ respectively. This indicates that nitrogen fertilizer application can increase grain yield potential at Estiewereda as compared to the other experimental locations on the other hand Fartawereda recorded lowest mean of grain yield. This result indicates that it has to be increase N-fertilizer application beyond 180 kg ha^-1^ in order to increase grain yield parallelism (Table 2&3) Highest grain yield was produced at 180 kg N ha^-1^ application for both malt barley cultivars, Holker and Miskal-21 that is 1361.1 and 1388.9kg ha^-1^, 2166.7and 2444.4 kg ha^-1^ and 1444.4 and 1722.2 kg ha^-1^ at Farta, Estie and Ebnat respectively. The present study shows that grain yield was significantly increased with increasing of N-fertilizer application. From this result we conclude that Miskal-21 was more responsive to N-fertilizer application as compared to Holker malt barley cultivar (Table 4, 5 & 6). This finding was in line with Mohammad*et al*., (2011) who observed that nitrogen application significantly affected grain yield of Barley varieties (local and sterling). Likewise Yadeta and Patricia (2012) reported that the cultivar and rate of N fertilizer interaction effect was also significant for grain yield. Therefore, this implies that the responses of two malt barley cultivars to rates of N fertilizer weredifferent. Miskal-21 had a linear and highly responsive to increased rate of N fertilizer for all locations (Table 4, 5 & 6). On the other hand, Holker had a linear but low responsive to increased rate of N-fertilizer for the tested locations (Table 4, 5 & 6).

**Table 3.**
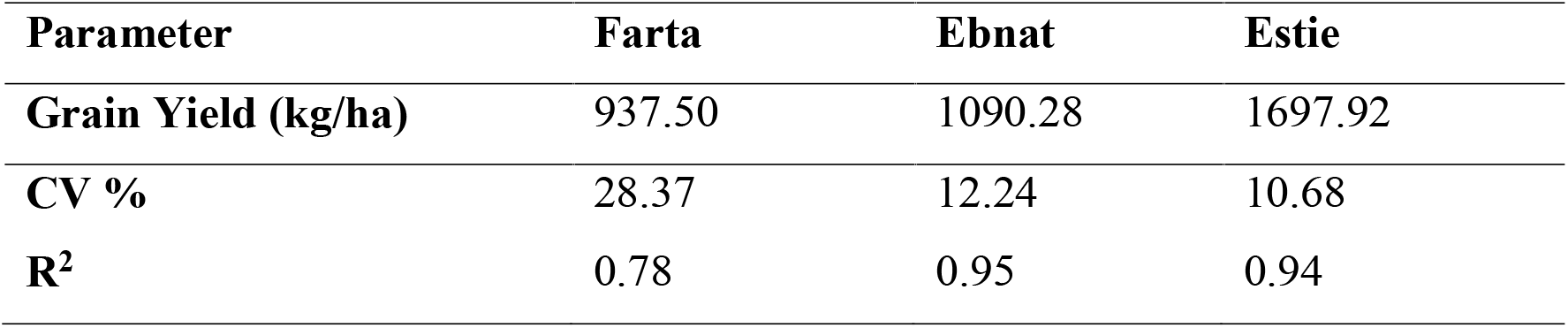
Environmental means of grain yield.

**Table 4.** Treatment mean of Grain yield of malt barley for Farta location.

| Parameter | N-fertilizer |  |  |  |  |  |  |  |
| --- | --- | --- | --- | --- | --- | --- | --- | --- |
|  | Holker |  |  |  | Miskal -21 |  |  |  |
|  | 0-N kg ha<sup>-1</sup> | 60-N kg ha<sup>-1</sup> | 120-N kg ha<sup>-1</sup> | 180-N kg ha<sup>-1</sup> | 0-N kg ha<sup>-1</sup> | 60-N kg ha<sup>-1</sup> | 120-N kg ha<sup>-1</sup> | 180-N kg ha<sup>-1</sup> |
| GY (kg/ha) | 555.6<sup>b</sup> | 805.6<sup>ab</sup> | 861.1<sup>ab</sup> | 1361.1<sup>a</sup> | 611.1<sup>b</sup> | 861.1<sup>ab</sup> | 1055.6<sup>ab</sup> | 1388.9<sup>a</sup> |

**Table 5.**
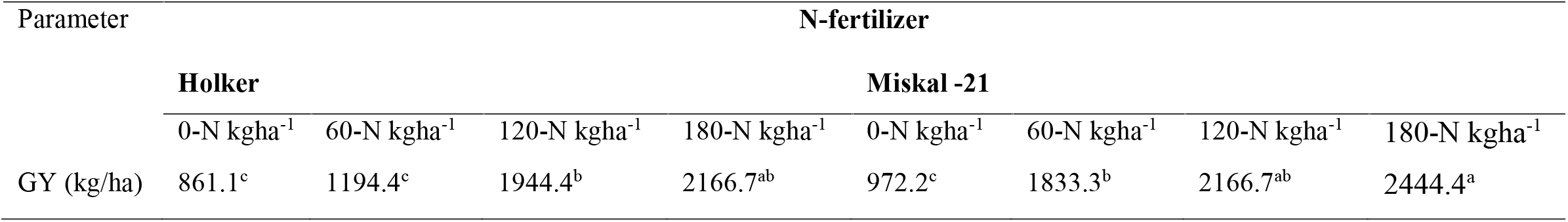
Treatment mean of Grain yield of malt barley for Estie location.

**Table 6.** Treatment mean of Grain yield of malt barley for Ebnat location.

| Parameter | N-fertilizer |  |  |  |  |  |  |  |
| --- | --- | --- | --- | --- | --- | --- | --- | --- |
|  | Holker |  |  |  | Miskal -21 |  |  |  |
| | 0-N kg$ha^{-1}$ | 60-N kg$ha^{-1}$ | 120-N kg$ha^{-1}$ | 180-N kg$ha^{-1}$ | 0-N kg$ha^{-1}$ | 60-N kg$ha^{-1}$ | 120-N kg$ha^{-1}$ | 180-N kg$ha^{-1}$ |
| GY (kg/ha) | 527.8<sup>d</sup> | 861.1<sup>c</sup> | 1111.1<sup>c</sup> | 1444.4<sup>ab</sup> | 416.7<sup>d</sup> | 1166.7<sup>bc</sup> | 1472.2<sup>ab</sup> | 1722.2<sup>a</sup> |

Nitrogen application had largely positive effects on grain yield production and productivity but negative effects on factors that would likely impact malting barley quality. From the present study it indicates that as increasing of nitrogen application there was also parallel increasing in grain yield in all the three weredas (Table 8& fig. 1.). Similarly, the study showed that increasing N-rates also enhancing the amount of protein content (Table 2& fig. 3).This positive correlation between grain yield and protein content (table 9 & 10) proven that both grain yield and protein content were highly dependent on the amount of nitrogen rates(fig. 1&fig. 3). From this resultwe can suggest that higher grain yield and protein content is advisable for food barley growers but in contrast with reasonable grain yield and lower protein content is recommended for malt barley growers.

**Figure 1.**
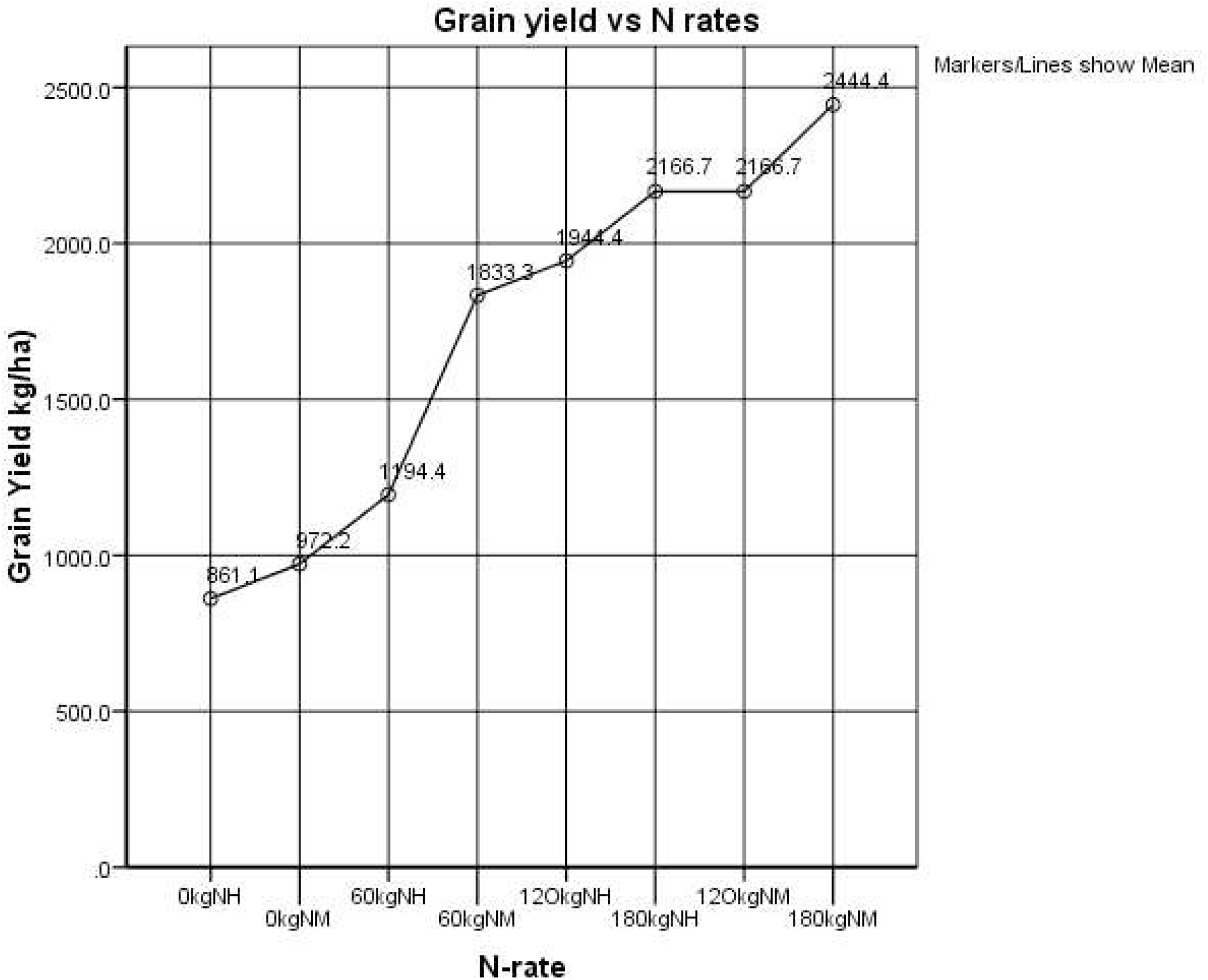
Relationship between mean grain yield of 2 malt barley variety and N rates

**Figure 2.**
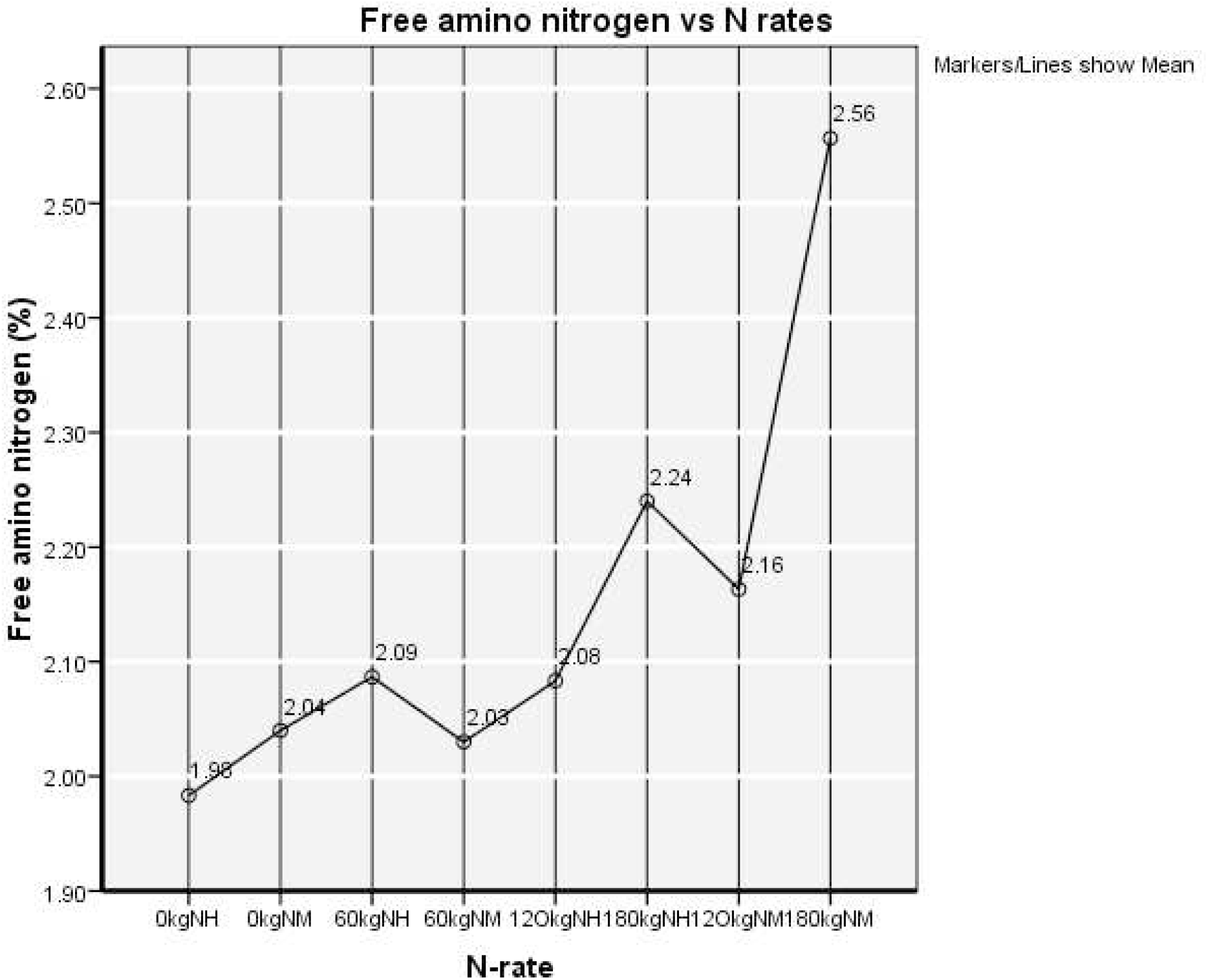
Relationship between mean free amino nitrogen of 2 malt barley variety and N rates

The present investigation indicated that highest grain yield was recorded at East-Estie as compare to the other weredas but similar protein content record was observed in all weredas at 180 kg/ha N (table 8). Holker was recorded lower grain yields as well as lower protein content as compared to miskal malt barley variety. Holker recorded 1444.4, 1361.1 and 2166.7kgha^-1^ grain yield and 14.33, 12.85 and 13.16% protein content at 180 kgha^-1^ N in Ebnat, Farta and Estie respectively. On the other hand, miskal recorded 1722.2, 1388.9 and 2444.6kgha^-1^ grain yield and 13.87, 14.39 and 14.32% in Ebnat, Farta and Estie (Table 8), respectively.

Therefore, the finding showed that yield increased up to 60, 120 and 180 kgha^-1^ N were a better N fertilization management that can optimized malt barley yield and malt qualities at Estie, Ebnat and Farta, respectively. The result recommended that malt breweries and malt growers must be used this rate of N at specified location in order to balance maximum yields with relatively low levels of protein. However, in view of the numerous negative effects of N fertilizer on malting barley quality outlined in this study, growers may be well-advised to limit N application to possibly 60 to 70% of the soil test recommendation in an effort to alleviate these negative effects. While this may result in reduced yield compared to applying N at higher rates, but economic returns is higher due to increased premium for malting compared to food barley.

### 4.2 Barley Protein

Lowest protein was observed at Farta experimental location and highest protein content was recorded at Estie (Table 7).As indicated in this result (Table 8), Miskal-21 was recorded highest protein content in all tested location. Hence this may results in lower extracts for the brewer. It also slows down water uptake during steeping, potentially affecting final malt quality. A very low protein level, on the other hand, resulted in a lack of enzymes necessary to modify the barley kernel and to break down the starch during brewing. In contrast to Miskal-21, Holker was relatively recorded lower protein content with reasonable yield reduction. The protein level of this malt variety was ranges from 11.5-13.8% as compared to Miskal-21 which was ranges from 12.5-14.9% (Table 8). Therefore the former indicated the suggested malt protein level that is best for brewers based on their process, their yeast and the type of beer they are making. This finding is agreed with the report of (Donovan *et al*., 2013) that they stated barley protein within the range of 11 – 12.5% can be used by maltsters to meet many brewers’ needs. They were also suggested protein level outside this range might be limited the requirement of maltster or brewers. Moreover, this result created a good insight that excessive rates of nitrogen were increased protein levels beyond the requirements of malt brewers. Therefore it is advisable to growers that application of nitrogen should be applied on the recommended rate in all location to obtain optimum yields as well as minor effect on the protein content (Fig.3).

**Table 7.** Environmental mean of malt barley yield and quality parameters.

**Tabale7. Environmental mean of malt barley yield and quality parameters**
| Parameters | Location |  |  |
| --- | --- | --- | --- |
|  | Ebnat | Estie | Farta |
| GY(kg/ha) | 1090.28 | 1697.92 | 937.50 |
| BP% | 12.97 | 13.08 | 12.53 |
| BM% | 8.81 | 11.14 | 9.72 |
| FAN | 156.4 | 138.7 | 167.6 |

**GY**- grain yield (kg/ha), **BP**-barley protein, **BM**-barley moisture and **FAN**-free amino nitrogen

**Table 8.** mean Effect of variety and N rate on yield and quality parameters.

**Tabale.8. mean Effect of variety and N rate on yield and quality parameters**
| Malt<br>barley<br>Variety | Location |  |  |  |  |  |  |  |  |  |  |  |  |
| --- | --- | --- | --- | --- | --- | --- | --- | --- | --- | --- | --- | --- | --- |
|  | Ebnat |  |  |  | Estie |  |  |  | Farta |  |  |  |  |
|  | N rate<br>(kg/ha) | GY(kg<br>/ha) | BP% | BM<br>% | FAN | GY(kg/<br>ha) | BP% | BM<br>% | FAN | GY(kg<br>/ha) | BP% | BM% | FAN |
| **Holker** | **0** | 527.8 | 12.45 | 9.42 | 2.01 | 861.1 | 12.61 | 10.70 | 1.98 | 555.6 | 13.4 | 9.19 | 2.27 |
|  | **60** | 861.1 | 13.13 | 8.84 | 2.23 | 1194.4 | 12.65 | 10.59 | 2.09 | 805.6 | 11.62 | 9.72 | 2.03 |
|  | **120** | 1111.1 | 13.40 | 9.51 | 1.84 | 1944.4 | 12.39 | 11.50 | 2.08 | 861.1 | 12.58 | 9.14 | 2.13 |
|  | **180** | 1444.4 | 14.33 | 9.17 | 2.30 | 2166.7 | 13.16 | 10.75 | 2.24 | 1361.1 | 12.85 | 10.46 | 2.02 |
| **Miskal** | **0** | 416.7 | 11.15 | 8.34 | 1.99 | 972.2 | 13.01 | 12.98 | 2.04 | 611.1 | 11.25 | 9.47 | 1.94 |
|  | **60** | 1166.7 | 12.28 | 8.88 | 2.10 | 1833.3 | 13.00 | 11.46 | 2.03 | 861.1 | 12.04 | 9.70 | 2.08 |
|  | **120** | 1472.2 | 13.11 | 8.06 | 2.18 | 2166.7 | 13.51 | 11.30 | 2.16 | 1055.6 | 12.35 | 10.24 | 2.19 |
|  | **180** | 1722.2 | 13.87 | 8.28 | 2.06 | 2444.6 | 14.32 | 9.84 | 2.56 | 1388.9 | 14.39 | 9.81 | 2.42 |

**GY**- grain yield (kg/ha), **BP**-barley protein, **BM**-barley moisture and **FAN**-free amino nitrogen

**Table 9.** Correlation coefficients among yield and quality traits of malt barley (Estie)

| Character | GY | FAN | BP | BM |
| --- | --- | --- | --- | --- |
| GY | 1 |  |  |  |
| FAN | 0.734 | 1 |  |  |
| BP | 0.633 | 0.871 | 1 |  |
| BM | -0.418 | -0.604 | -0.387 | 1 |

\*significant at the 0.05 level, \*\* significant at the 0.01 level, **GY**- grain yield (kg/ha), **BP**-barley protein, **BM**-barley moisture and **FAN**-free amino nitrogen

**Table 10.**
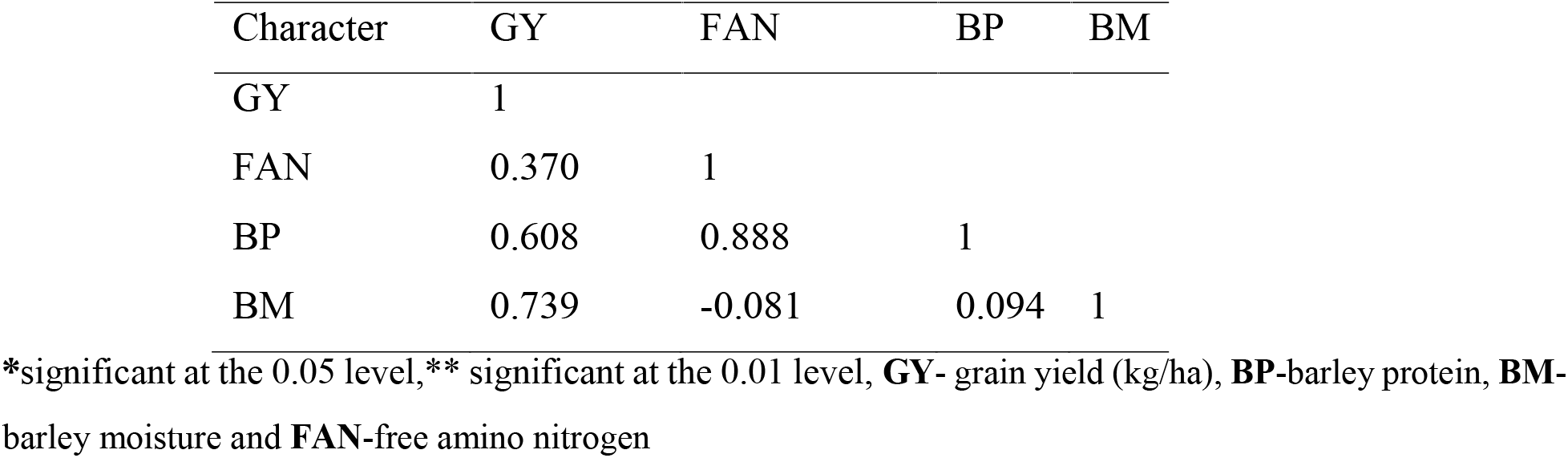
Correlation coefficients among yield and quality traits of malt barley (Farta)

| Character | GY | FAN | BP | BM |
| --- | --- | --- | --- | --- |
| GY | 1 |  |  |  |
| FAN | 0.370 | 1 |  |  |
| BP | 0.608 | 0.888 | 1 |  |
| BM | 0.739 | -0.081 | 0.094 | 1 |

\*significant at the 0.05 level, \*\* significant at the 0.01 level, **GY**- grain yield (kg/ha), **BP**-barley protein, **BM**-barley moisture and **FAN**-free amino nitrogen

**Figure 3.**
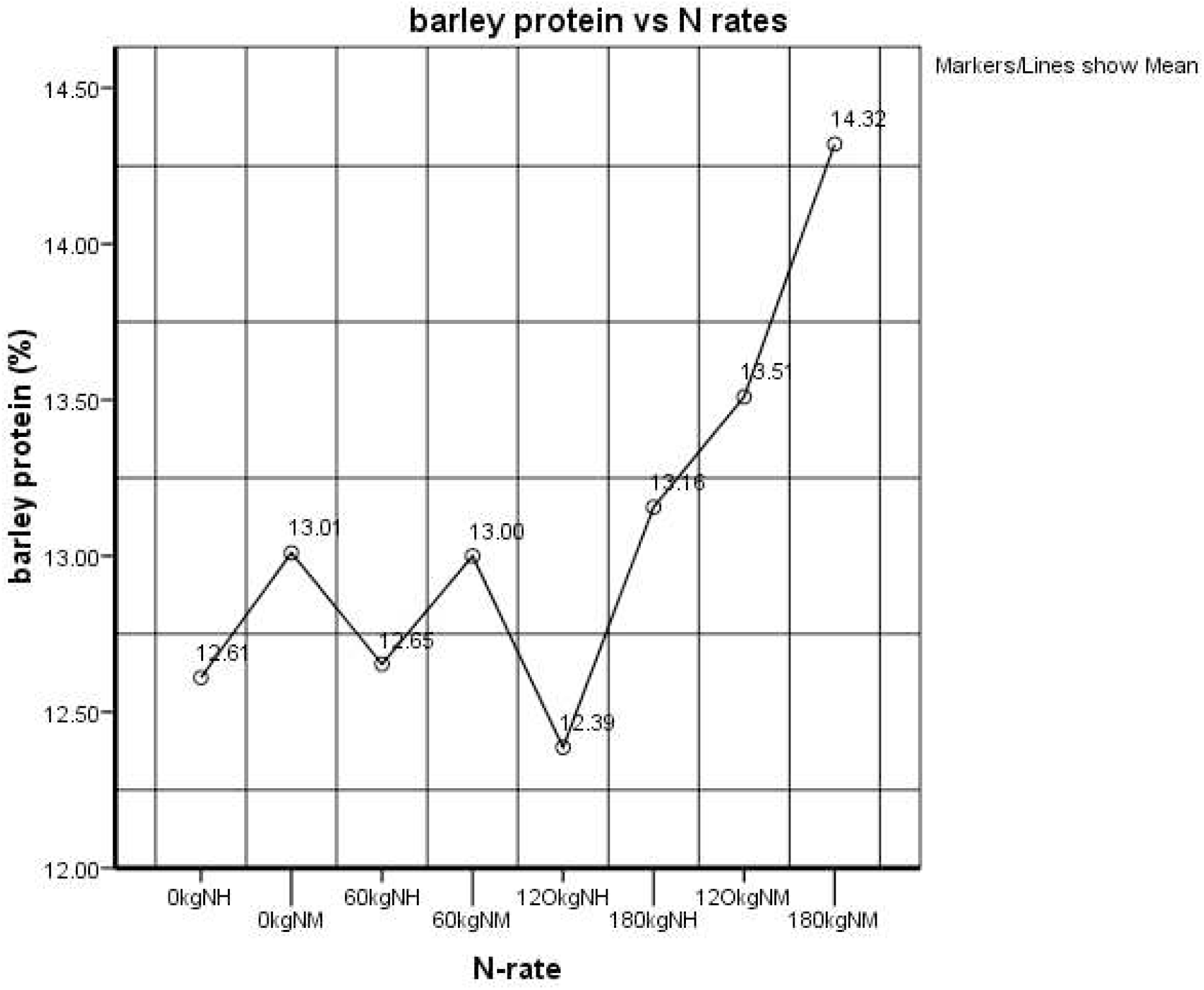
Relationship between mean barley protein percentage of 2 malt barley variety and N rates

### 4.3. Barley Moisture Content

Excellent moisture content was recorded at Ebnat, Farta and Estie respectively. This result indicated a good record moisture content in all locations hence Malting barley over 13.5% moisture does not store well and can significantly reduce the malt qualities. From the present finding, the mean barley moisture content ranged from 8.81-11.14 across locations. This moisture is low enough to inactivate the enzymes involved in seed germination as well as to prevent heat damage and the growth of disease microorganisms. Therefore, quality and germination capacity may significantly deteriorate and growers should be advised to maintain barley moisture below 13.5% when store.

## REFERENCE

AACC (American Association of Cereal Chemists) 2000. Approved Methods of the American Association of Cereal Chemists. 10th ed., Amercan Association of Cereal Chemists, St Paul, MN.

Aaron L. M., Michael J. E. and Marta S. I. 2012. Quality of western Canadian malting barley– Canadian Grain Commission

Abeledo L.G., Calderini D.F., and Slafer G.A. 2003. Genetic improvement of yield responsiveness to nitrogen fertilization and its physiological determinants in barley. Euphytica, 133: 291–298.

Acuna M.L, Savin R, Cura J.L, Slafer G. A 2005. Grain protein quality in response to changes in pre-anthesis duration in wheats released in1940, 1964 and 1994. J. Agron. Crop Sci. 191: 226–232.

AOAC (Association of Official Agricultural Chemists) 1990. Official Methods of Analysis. Arlington, Virginia, USA

Donovan J.T., Turkington T.K., Edney E., Smith E. and Harker K. N. (2013). Varietal and agronomic impacts on malt barley yield and quality. Farm tech 2013 conference proceeding.

FAO (Food and Agricultural Organization). 2009. Malt barley beer. Rome, Italy

Feiten Abay, Edema, R. and Bjornstad A. 2010. Development of improved scald tolerant barley varieties with superior endues (malt) and nutritional quality for dry land areas of the northern Ethiopia. Second Ruforum. Biennial Meeting 20 - 24 September, Entebbe, Uganda

Fekadu A, Berhanu B, Fekadu F, Adisie N, Tesfaye G. 1996. Malting Barley Breeding. pp. 24-33. Proceedings of the First Barley Research Review Workshop, 16 to 19 October 1993. Addis Ababa: IAR/ICARDA

Jürg M. B., David D. B., Kenneth G. C., Stephen C. M. and Alexander D. P. 2008. Importance and Effect of Nitrogen on Crop Quality and Health. Elsevier Inc. pp 51–70

Leistrumaite A, and Paplauskiene V 2005. Genetic resources of spring barley: screening for yield stability and grain malt quality traits. Biologija, 3: 23–26.

McKenzie R.H., Middleton A.B., and Bremer E 2005. Fertilization, seeding date, and seeding rate for malting barley yield and quality in southern Alberta. Can. J. Plant Sci. 85: 603–614.

Mengel K, Hutsch B, and Kane Y 2006. Nitrogen fertilizer application rates on cereal crops according to available mineral and organic soil nitrogen. Eur. J. Agron. 24: 343–348.

Mohammed, H. and Getachew L 2003. An Overview of malt barley production and marketing in Arsi. pp. 1–25. Proceedings of the Workshop on Constraints and Prospects of Malt Barley, Production, Supply, and Marketing Organized by Asella Malt Factory and Industrial Projects Service. March 15, 2003.

Palmer G.H 2000. Malt performance is more related to inhomogeneity of protein and b-glucan breakdown than to standard malt analyses. J. Inst. Brew. 106: 189–192.

Pettersson C.G. 2007. Predicting Malting Barley Protein Concentration based on canopy reflectance and site characteristics. [PhD. Thesis]. Swedish University of Agricultural Sciences. SE-750 07 Uppsala, Sweden.

Pržulj N, Momčilović V, Nožinić M, Jestrović Z, Pavlović M,and Orbović B. 2010. Importance and breeding of barley and oats. Field Veg. Crop. Res. 47(1): 33–42

Snežana J., Đorđe G., Radojka M. S., Nikola H. and Jela I. 2011. Effects of nitrogen fertilization on yield and grain quality in malting barley. African Journal of Biotechnology 10.:19534–19541

Teshome Galano, Geremew Bultosa and Chemeda Fininsa. 2011. Malt quality of 4 barley (Hordeum vulgare L.) grain varieties grown under low severity of net blotch at holetta west shewa, Ethiopia. Africa Journal of Biotechnology. 10:797–806

Wang J.M., Zhang G.P, and Chen J.X. 2001. Cultivar and environmental effects on protein content and grain weight of malting barley. J. Zhejiang University (Agricultural and Life Sciences) 27: 503–507.

